# ADULT EXPRESSION OF SEMAPHORINS AND PLEXINS IS ESSENTIAL FOR MOTONEURON SURVIVAL

**DOI:** 10.1101/2022.10.30.514429

**Authors:** Aarya Vaikakkara Chithran, Douglas W. Allan, Timothy P. O’Connor

## Abstract

A role for axon guidance genes in the adult nervous system has not been fully elucidated. We performed an RNAi screen against guidance genes in the adult *Drosophila melanogaster* nervous system and identified fourteen genes required for adult survival and normal motility. Additionally, we show that adult expression of Semaphorins and Plexins in motoneurons is necessary for neuronal survival, indicating that guidance genes have critical functions in the mature nervous system.

## Introduction

An extraordinary characteristic of the nervous system is the complexity of its wiring. Although there are multiple mechanisms involved in establishing these connections, a key event is the guidance of axons to their specific targets. During the development of the nervous system, axons navigate to their targets in response to various attractive and repulsive guidance cues^1^. Recently, it has become clear that after functional circuits have been established, many neurons continue to express developmental guidance genes. Indeed, gene expression studies in the adult rodent brain have identified brain regions that are enriched for genes involved in neuronal development and axon guidance^2^.

The expression of guidance genes in the adult indicates that there are likely additional roles for them beyond the initial phase of axonal outgrowth and synaptogenesis^3^. Indeed, blocking the function of class3 Semaphorins and Ephrins promotes axonal sprouting and regeneration after nerve injury^4-7^. Guidance cues are also known to regulate adult synaptic plasticity. Conditional deletion of the receptor for Netrin-1 (DCC) in adult mice forebrain neurons results in shorter dendritic spines, loss of long-term potentiation, and impaired spatial and recognition memory^8^. Similarly, Semaphorin 5B regulates the elimination of synaptic connections in cultured hippocampal neurons^9^. Axon guidance genes have also been linked to neurodegenerative diseases. Given their role in axonal guidance and synapse regulation, it has been proposed that changes in guidance protein expression or function may trigger defects in neuronal networks leading to neuronal dysfunction and loss^10^. Thus, based on their known functions during development, their expression in the adult brain and possible implications in neurodegenerative diseases, we examined whether guidance genes are critical for the maintenance and survival of neurons in the mature nervous system.

## Results

### Knockdown of a subset of axon guidance genes reduces adult survival

Using the expression pattern data^11^ available on FlyBase^12-13^, we compared the expression of axon guidance genes in the embryonic (10-24 hours) and adult (1-20 days) *Drosophila melanogaster*. We found that more than 96% of the guidance genes expressed in the embryo continue to be expressed in adults. Query results from the RNA-Seq Expression Profile search tool showed that 162 genes expressed in the adult *Drosophila* nervous system were categorized under the biological process Gene Ontology (GO) term ‘axon guidance’ and 122 genes under ‘dendrite morphogenesis’ (Supplementary Tables 1, 2). After excluding transcription factors, 44 genes were prioritized based on their previously known roles in axonal guidance and the availability of genetic tools (Supplementary Tables 3, 4). We refer to these genes collectively as ‘axon guidance genes’. Based on the molecular function GO terms provided on FlyBase, these 44 axon guidance genes were categorized as ‘actin/microtubule-binding’, ‘cell adhesion’, ‘guidance ligand/protein binding’ and ‘receptor activity’ (Figure. 1).

**Figure 1.**
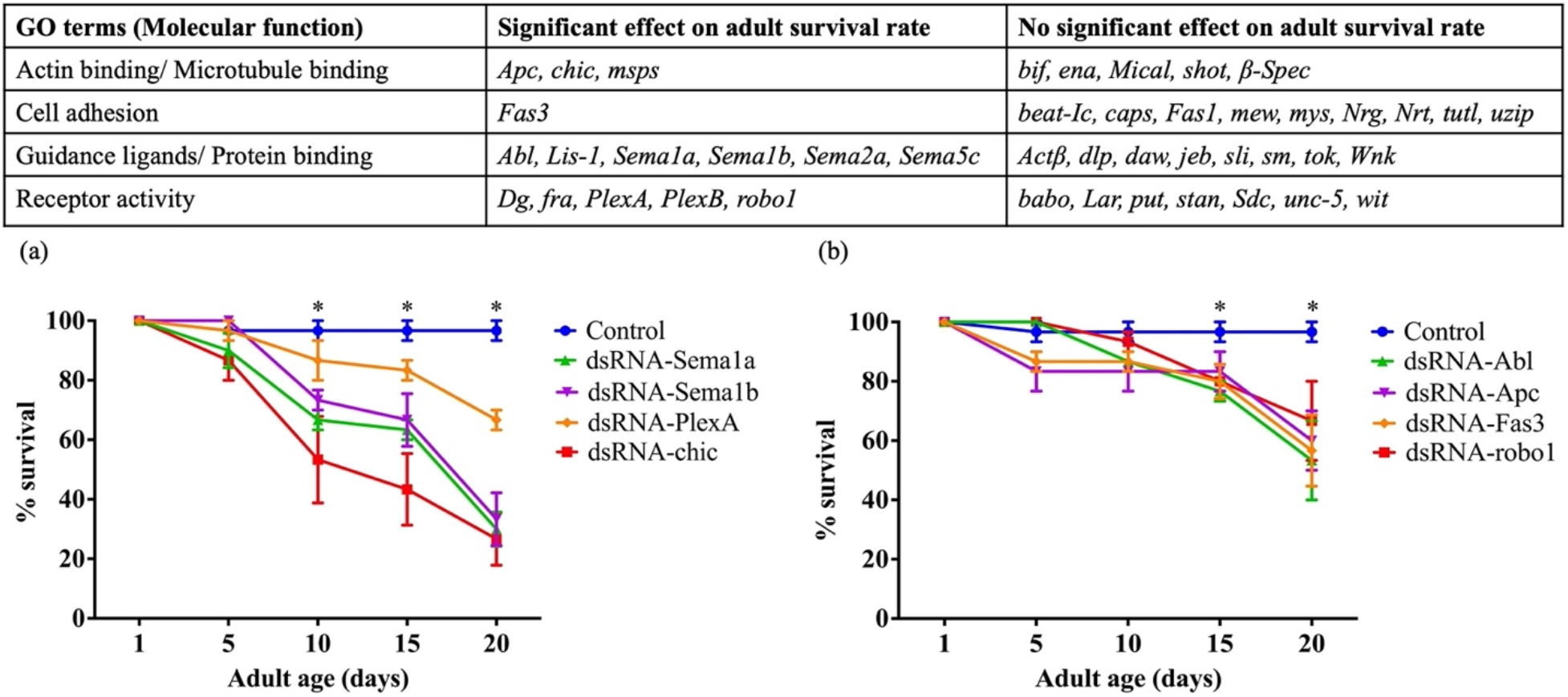
Knockdown of a subset of axon guidance genes reduces adult survival rate. The table summarizes the axon guidance genes that were tested for the impact of knockdown on adult survival rate. The axon guidance cues are categorized into four groups based on the molecular function GO terms. (a) Survival analysis curves of examples of genes that showed a significant effect on survival from day 9 post eclosion. (b) Survival analysis curves of genes that showed a significant effect from day 14. Data shown are mean ± SEM (p<0.05, two-way ANOVA and Tukey’s multiple comparison tests). ‘*’ indicates the timepoints when the survival of knockdowns was significantly different from that of the controls.

In an initial screen to identify whether any genes were essential for viability, we knocked down the expression of all 44 genes (using multiple *dsRNA* and *shRNA* lines; see Methods Table 2) selectively in the adult nervous system, using the GeneSwitch (*elav-GeneSwitch-Gal4*) and TARGET (*elav-Gal4* with *tub-Gal80^ts^*) systems^14^ (see Methods Table 1). In the GeneSwitch screen, knockdown of 15 of the 44 genes showed significant reduction in adult survival rate. The TARGET system screen confirmed these data for 14 of the 15 genes. (Figure 1, Supplementary Figure 1). In control flies, 97% survive to day 20 post eclosion, whereas the percentage survival post knockdown for 14 genes were significantly reduced to 27% (*chic*), 30% (*Sema1a*), 33% (*fra, Sema1b*), 37% (*Sema5c*), 40% (*PlexB*), 53% (*Abl, Dg, msps*), 57% (*Fas3*), 60% (*Apc*), 67% (*PlexA, robo1*), and 77% (*Sema2a*).

**Table 1.**
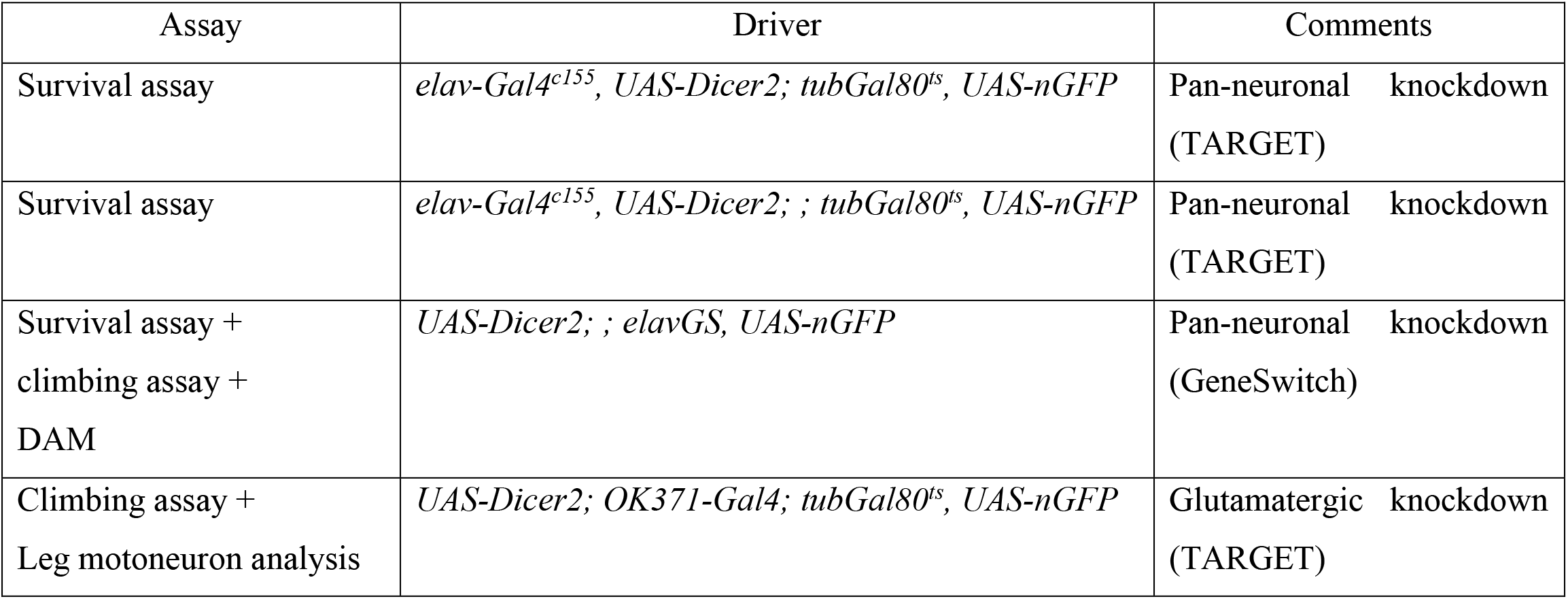
List of Gal4 driver lines used in the study.

**Table 2.**
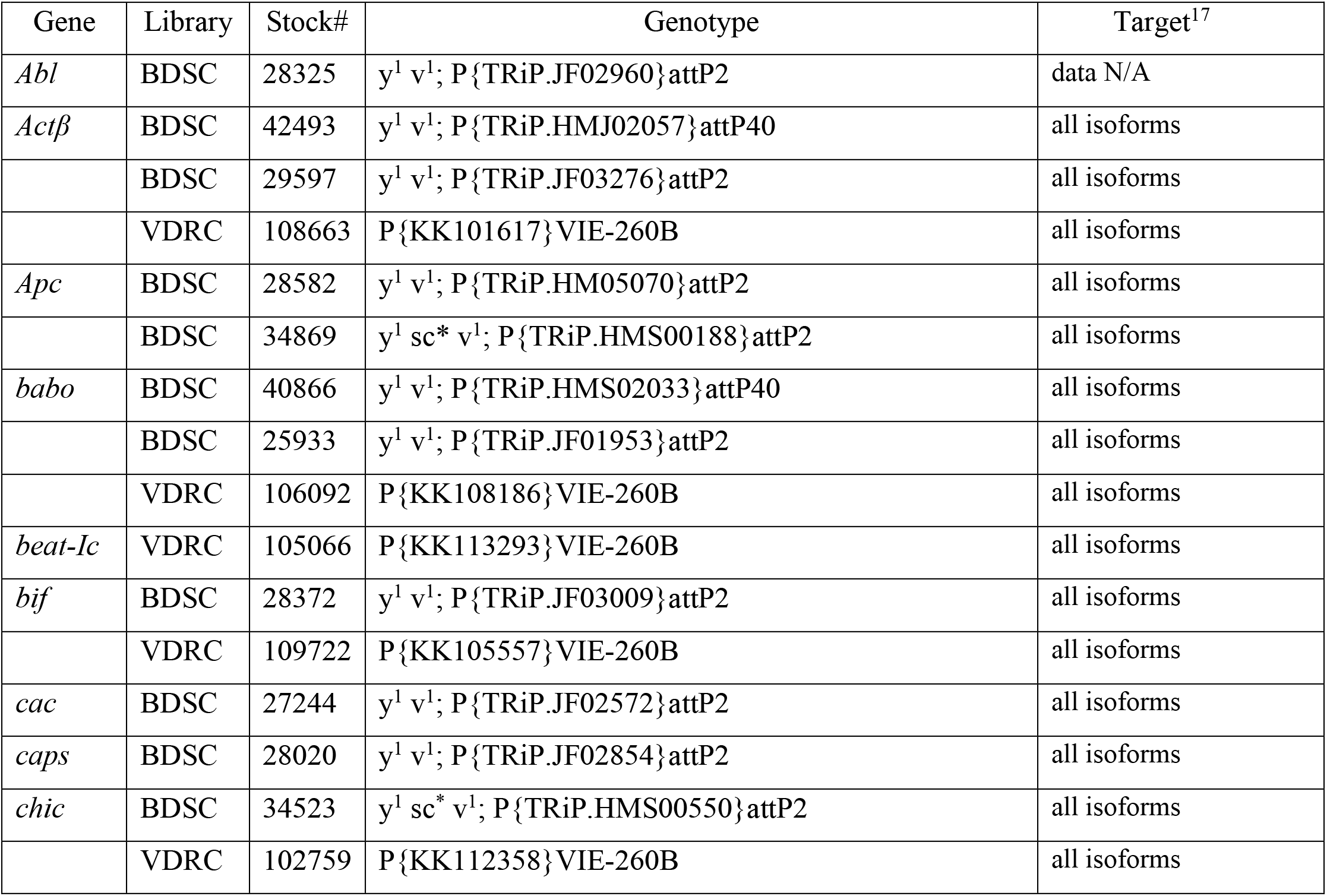
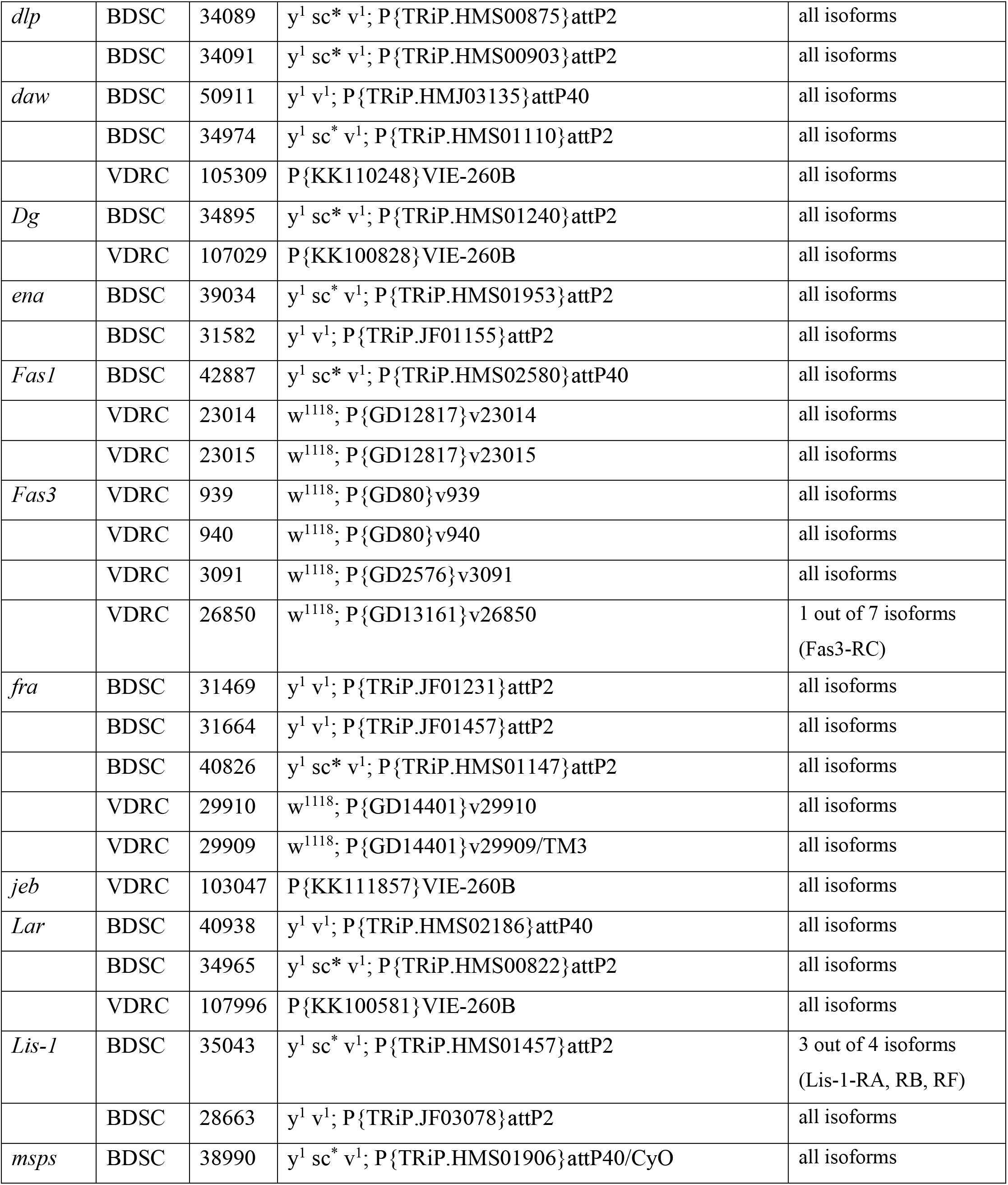
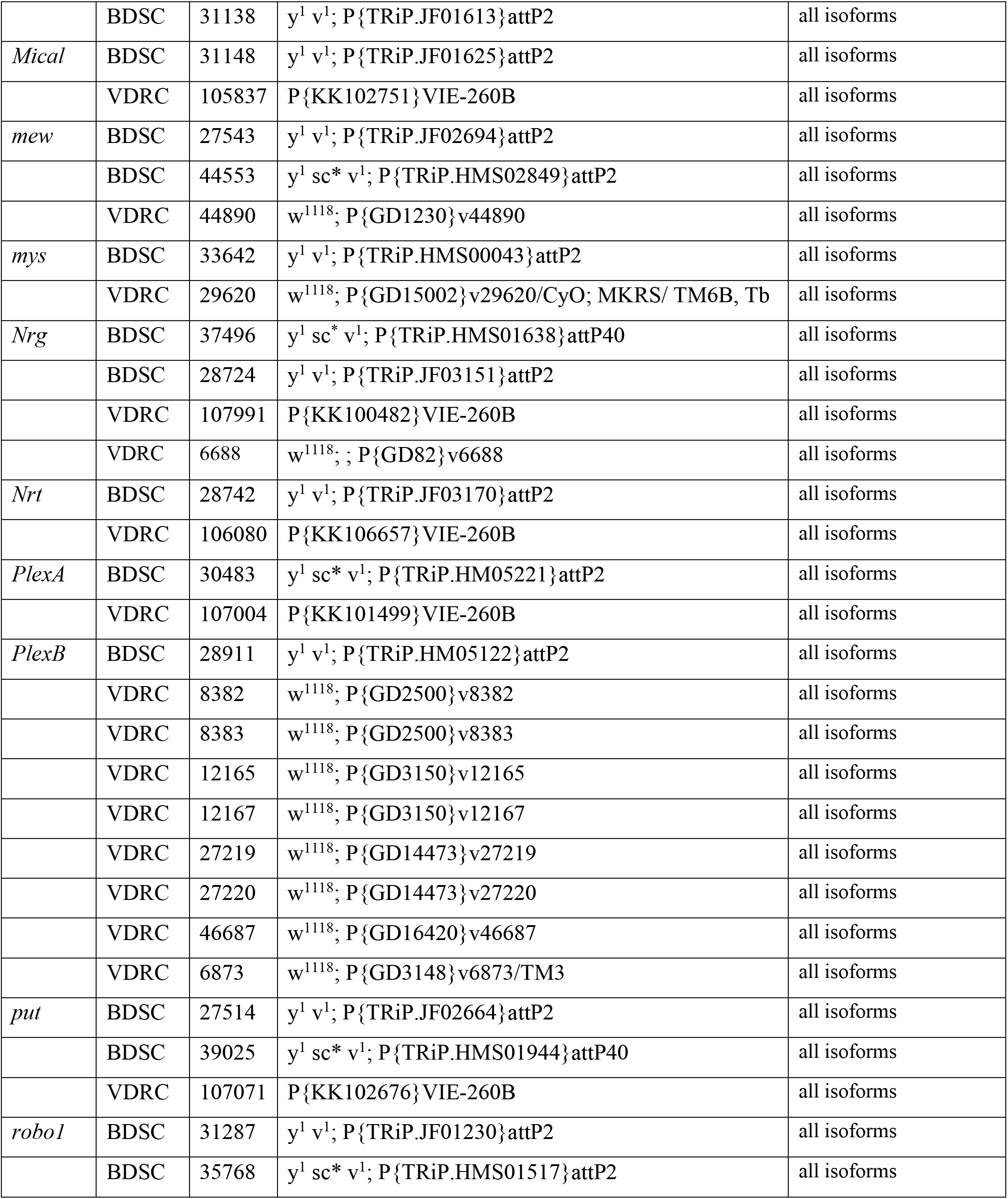
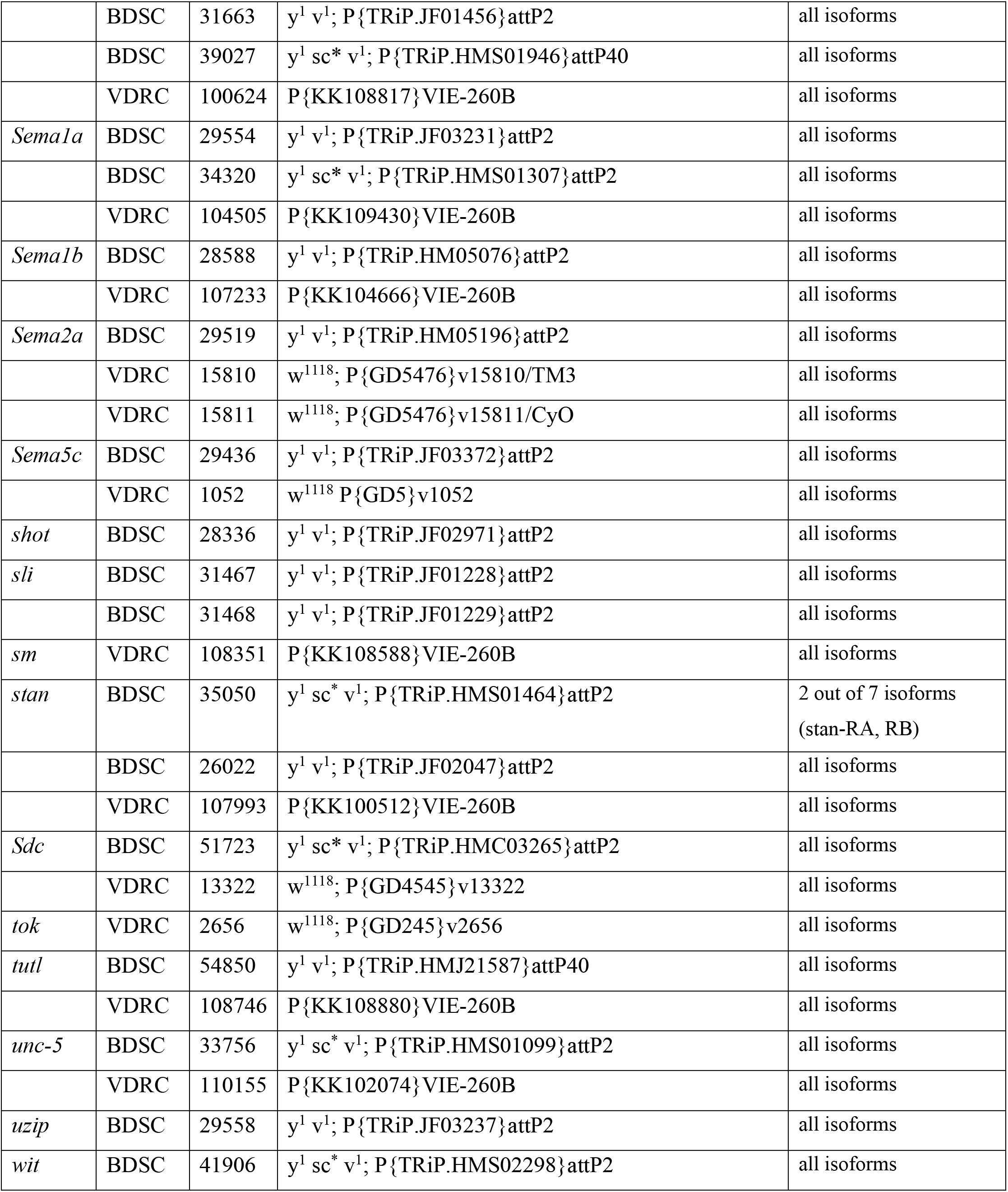
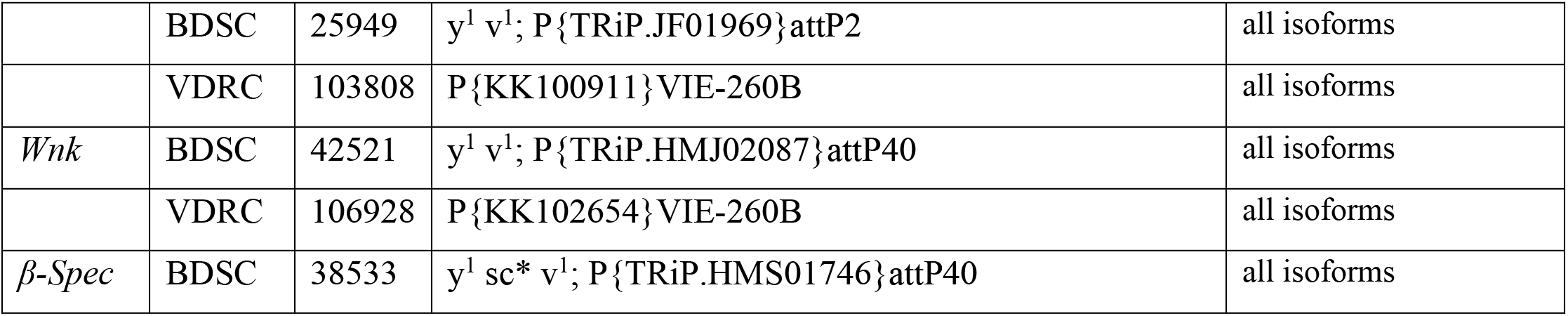
List of *UAS-shRNA* and -*dsRNA* lines used in the study.

### Motility is impacted by the knockdown of specific axon guidance genes

To examine adult fly behavior in the period before survival declines, we focused on these 14 genes and divided them into two groups based on their survival analysis graphs: 1) Those that started to show reduction in adult survival from approximately day 9 onwards (*Sema1a, Sema1b, Sema2a, Sema5c, PlexA, PlexB, chic*), and 2) Those that started to show reduction from approximately day 14 onwards (*Abl, Apc, Dg, Fas3, fra, msps, robo1*). Using these two time points, we conducted motility assays with group one at 9 days and group two at 14 days, after the initiation of *shRNA* and *dsRNA* expression. Due to potential differences in the effectiveness and timing of knockdown of various *UAS-shRNA* and *-dsRNA* lines, each knockdown was compared only to their respective controls (see Methods Table 3). As an initial examination of motility, we tested whether panneuronal knockdown (*elav-GeneSwitch-Gal4*) of the genes impacted adult climbing behavior. We found that flies that had axon guidance genes knocked down showed a significant climbing defect. Flies that failed to reach the target line were either very slow or did not start climbing even after tapping the vials. In the control flies, success of climbing ranged from 87% to 94%. We observed the following climbing success rates in knockdowns of *Sema1a* (72%), *Sema2a* (63%), *PlexA* (38%) and *PlexB* (37%). The climbing success rates ranged from 10% to 59% in the knockdowns of the other 10 genes (Figure 2a, Supplementary Figure 2a).

**Table 3.** List of control lines used in the study.

|  |  |  |  |
| --- | --- | --- | --- |
| attP2 insertion TRiP line injection strain | BDSC | 36303 | y<sup>1</sup> v<sup>1</sup>; P{CaryP}attP2 |
| attP40 insertion TRiP line injection strain | BDSC | 36304 | y<sup>1</sup> v<sup>1</sup>; P{CaryP}attP40 |
| KK line injection strain | VDRC | 60100 | y,w<sup>1118</sup>; P{attP,y<sup>+</sup>,w[3']} |
| GD line injection strain | VDRC | 60000 | w<sup>1118</sup> |
| UAS-dsRNA-GFP on 2<sup>nd</sup> chromosome | BDSC | 9331 | w<sup>1118</sup>; P{UAS-GFP.dsRNA.R}143 |
| UAS-dsRNA-GFP on 3<sup>rd</sup> chromosome | BDSC | 9330 | w<sup>1118</sup>; P{UAS-GFP.dsRNA.R}142 |
| UAS-dsRNA-mCherry (additional control for VALIUM 20 attP2 insertion lines) | BDSC | 35785 | y1 sc* v1; P{VALIUM20-mCherry}attP2 |
| UAS-mCherry (additional control for VALIUM 10 and VALIUM 20 lines) | BDSC | 35787 | y1 sc* v1; P{VALIUM10-mCherry}attP2 |
| UAS-dsRNA-lacZ (additional control for KK lines) | VDRC | 51446 | w<sup>1118</sup>; P{GD936}v51446 |

**Figure 2.**
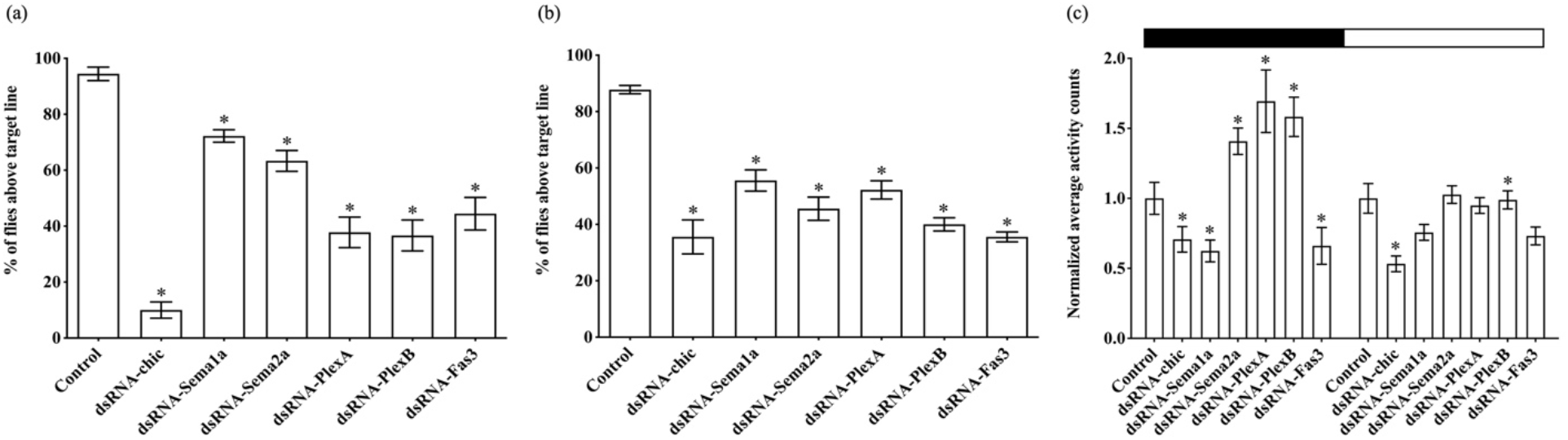
Motility is impacted by the knockdown of specific axon guidance genes. (a) Effect of pan-neuronal knockdown of *chic, Sema1a, Sema2a, PlexA, PlexB* and *Fas3* (using *elav-GeneSwitch-Gal4*) on adult climbing behavior. A significant decrease in adult climbing ability was observed after knockdown. (b) Similar observations were made when each gene was knocked down specifically in glutamatergic motoneurons using the *OK371-Gal4*. Data shown are mean ± SEM (p<0.05, one-way ANOVA and Tukey’s multiple comparison tests). (c) Effect of panneuronal knockdown of axon guidance genes (using *elav-GeneSwitch-Gal4*) on adult activity (measured using DAM). The bar graph compares the normalized average activity counts per 12 hours for each gene from multiple flies over a period of 3 days in 12 hours light and 12 hours dark cycle. The black portion (0-12 hours) represents the dark cycle, and the white portion (12-24 hours) represents the light cycle. Data shown are mean ± SEM (‘*’ denotes p<0.05, one-way ANOVA and Tukey’s multiple comparison tests).

Since a significant decline in adult climbing behavior was observed after pan-neuronal knockdown, we tested whether knocking down genes primarily in motoneurons would induce a similar phenotype using a glutamatergic neuron specific Gal4 driver (*OK371-Gal4*). Again, flies with knocked down guidance genes showed defects in adult climbing behavior (Figure 2b, Supplementary Figure 2b). Comparing the severity of climbing defects caused by the knockdown of each gene using the two drivers, only *chic* and *Apc* showed less severity after knockdown in motoneurons compared to all neurons. The remaining 12 genes showed no significant difference in the severity of climbing phenotype between the two Gal4 drivers (Figure 2a-b, Supplementary Figure 2a-b).

We further examined the motility of adult flies using a *Drosophila* Activity Monitor (DAM) by continuously monitoring individual flies for 3 days. *Fas3* and *chic* knockdown flies showed significantly lower overall activity counts than the controls throughout the day (both light and dark cycles). *Sema1a* knockdown flies exhibited significantly lower activity counts in both cycles whereas *Sema2a, PlexA* and *PlexB* knockdown flies exhibited significantly higher activity counts during the dark cycle (Figure 2c). Flies also showed aberrant activity counts following knockdown of *Abl, msps* and *robo1* (Supplementary Figure 2c).

### Knockdown of Semaphorins and Plexins results in motoneuron death

To investigate why motility was impacted after knockdown, we examined the number of leg motoneurons in the thoracic clusters of adult ventral nerve cords in a subset of the 14 genes using the *OK371-Gal4* driver (Figure 3a). Knockdown of *Sema1a, Sema2a, PlexA* and *PlexB* resulted in a significant loss of motoneurons in each thoracic cluster (T1, T2, T3). In T1, total neurons were reduced by 32%, 33%, 30% and 38% in *Sema1a, Sema2a, PlexA* and *PlexB* knockdowns respectively. The following reductions were observed in T2 and T3 segments respectively: 44% and 42% (*Sema1a*), 41% and 49% (*Sema2a*), 32% and 36% (*PlexA*) and 30% and 34% (*PlexB*) (Figure 3b-e, Supplementary Figure 3).

**Figure 3.**
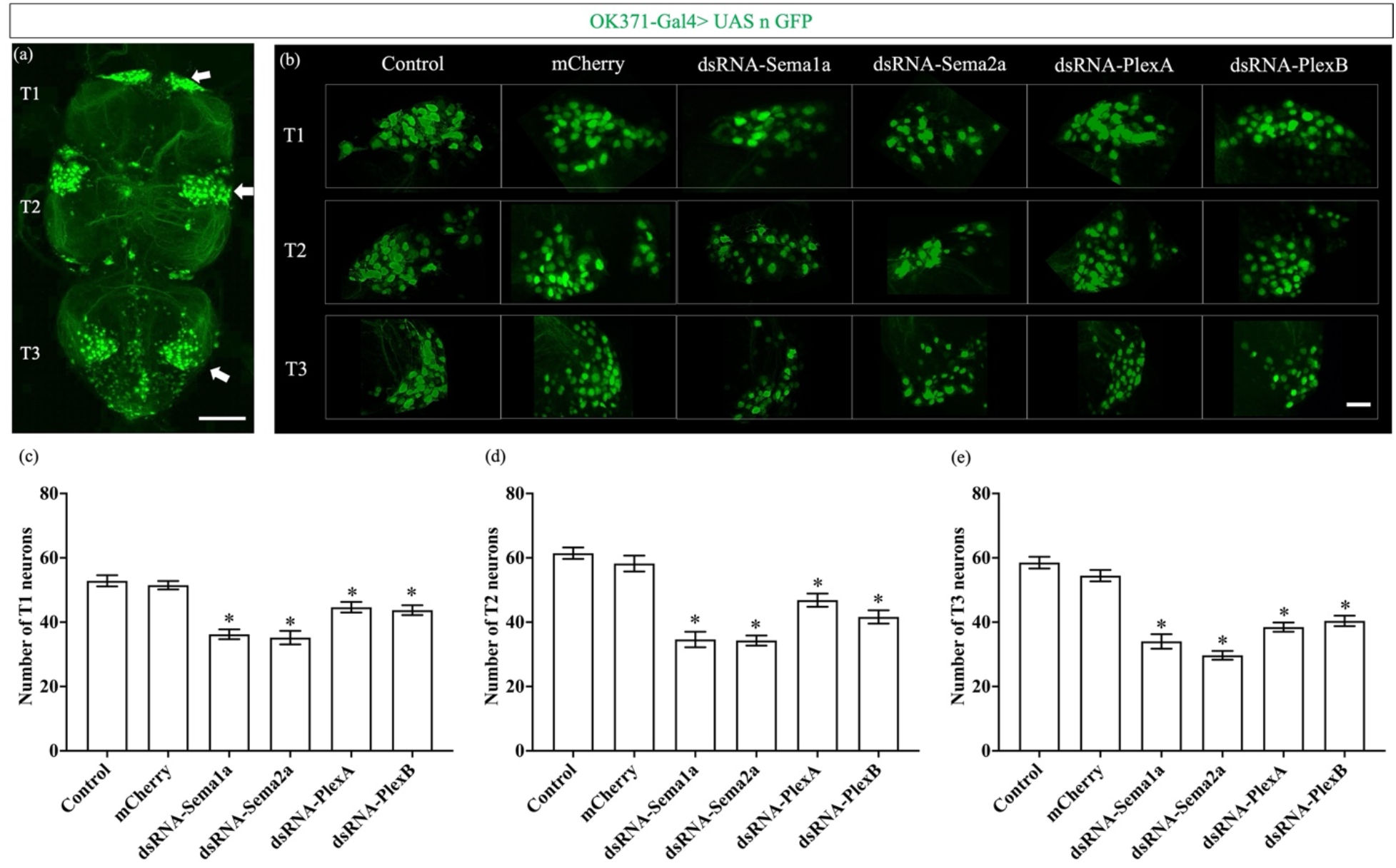
Knockdown of Semaphorins and Plexins results in motoneuron death. (a) Adult ventral nerve cord showing neuronal GFP expression driven by *OK371-Gal4*. White arrows point to the location of leg motoneuron cell bodies in T1, T2, T3 thoracic segments. Scale bar represents 100 μm. (b) Effect of glutamatergic knockdown of Semaphorins and Plexins (using *OK371-Gal4*) on adult leg motoneuron survival. A decrease in the number of leg motoneurons was observed after knockdown of Semaphorins and Plexins genes. Scale bar represents 20 μm. (c), (d), (e) Quantification of adult leg motoneurons in T1, T2, and T3 respectively, following 9 days of axon guidance gene knockdown. Data shown are mean ± SEM (‘*’ denotes p<0.05, one-way ANOVA and Tukey’s multiple comparison tests, n=20).

## Discussion

Among the 44 axon guidance genes we screened, 14 genes were essential for adult survival. Mutations in these 14 axon guidance genes were previously known to cause lethality during development (lethal alleles listed on FlyBase^12^ under FBcv:0000351). This implies that axon guidance genes are essential throughout the lifespan of *Drosophila,* from development through the adult. To better understand the cellular impact after guidance gene knockdown, we used motility assays to focus our examination on specific neural circuits. Employing a climbing assay and the DAM system, we found a significant impact on motility in knockdown flies. Since knockdown was restricted to adult flies, these locomotor phenotypes were not the result of developmental defects but are likely due to defects in the adult motoneuron circuit. This was confirmed when knockdown of *Sema1a, Sema2a, PlexA* and *PlexB* resulted in a significant loss of motoneurons in all three thoracic clusters (T1, T2, T3). These observations suggest that axon guidance proteins may act cell autonomously within motoneurons or with neighboring motoneurons to maintain their survival. Further studies on Semaphorin signaling downstream events will be required to better understand the mechanism underlying this phenotype.

The main goal of our study was to examine whether axon guidance genes have an important function in the adult. We identified 14 axon guidance genes that are essential for survival and normal motility. From our database search, we found that many of the embryonic axon guidance genes continue to be expressed in the adult. Similarly, gene expression studies in adult mouse brain suggest that although the expression patterns of many genes change dramatically during development, the brain retains a degree of the embryological gene expression that is important for the maintenance of established units in the adult brain^15^. Studies in rhesus monkeys (*Macaca mulatta*) have identified sets of genes that are important equally for the formation as well as the maintenance and plasticity of connections in the adult thalamus^16^. These imply that the expression of axon guidance genes in the adult are not just a “residue” of the developmental pattern that reflects processes occurring when connectivity is established, but are functionally relevant in adults. Indeed, our results show that expression of axon guidance genes in adulthood is required for organism viability and neuronal survival.

## Methods

### Prioritizing axon guidance genes for the screen

The high throughput expression pattern data^11^ accessed through the RNA-Seq Expression Profile search tool from FlyBase^12-13^ (www.flybase.org) version FB2013_05 (released September, 2013) was used to search for genes expressed in *Drosophila melanogaster* in the embryo (stage-wise expression, 10-24 hours) and in the adult (tissue-wise expression in head, 1-20 days post eclosion). An ‘expression on’ filter was used to search for genes expressed above a certain expression profile threshold, namely very low (1-3 RPKM), low (4-10 RPKM), moderate (11-25 RPKM), moderately high (26-50 RPKM), high (51-100 RPKM), very high (101-1000 RPKM) and extremely high (>1000 RPKM). The resulting dataset was further refined using biological process GO terms ‘axon guidance’ and ‘dendrite morphogenesis’. Transcription factors were excluded using cellular component GO term ‘nucleus’. Gene list Venn diagram (http://genevenn.sourceforge.net/) was used to categorize the genes into discrete expression levels, ranging from very low to extremely high, and to compare the genes expressed in the embryo and adult *Drosophila*. These data combined with their previously known roles in receptor activity and signalling during neuronal pathfinding, and the availability of genetic tools were used to prioritize genes.

### Fly genetics

Reducing axon guidance gene expression by RNAi in the adult nervous system was carried out using two well-characterized systems: 1) GeneSwitch and 2) TARGET. Gal4 driver line virgin females (see Table 1 below) were crossed with *UAS-shRNA/dsRNA* males (see Table 2 below) or the appropriate negative control line males (see Table 3 below). In case of the GeneSwitch screen, crosses were set up at 25°C. The progeny (F1) was fed with food containing 6.5μg/mL mifepristone (RU486) from day 1 post eclosion to activate the Gal4-UAS system. In case of the TARGET screen, crosses were set up at 18°C and the progeny (F1) was switched to 29°C immediately after eclosion to repress Gal80^ts^ and activate the Gal4-UAS system.

### Fly stocks

*Drosophila melanogaster* stocks used in this study were maintained on standard cornmeal food and at 18, 25, or 29°C in environment rooms set at 70% humidity. The stocks used in this study are as follows:

### Survival assay

F1 progeny males were collected immediately after eclosion. They were maintained in vials of 10 and their survival was recorded every day for the next 20 days. Flies were flipped into fresh vials every week. Survival analysis for the GeneSwitch screen was carried out at 25°C and that for the TARGET screen was carried out at 29°C. In both cases, multiple *UAS-dsRNA/shRNA* lines were tested for each axon guidance gene. For each line tested, 3 vials containing 10 flies were assayed. The results were analysed using two-way ANOVA and Tukey’s multiple comparison tests using GraphPad Prism.

### Climbing assay

To identify adult climbing defects, F1 male flies were collected immediately after eclosion, maintained in groups of 10 flies per vial and assayed at a desired age. During the climbing assay, flies were transferred to a clean, empty vial with a line drawn 7.5 cm from the bottom. Flies were allowed to acclimatize for 1 minute and then tapped to the bottom to induce an innate climbing response. The number of flies that successfully reached the 7.5 cm line in 8 seconds were recorded. For each genotype, 3 vials were assayed and 3 replicates per vial were performed to ensure an accurate reading for each vial. The results were analysed using one-way ANOVA and Tukey’s multiple comparison tests using GraphPad Prism.

### *Drosophila* Activity Monitor (DAM) System

The DAM system from TriKinetics was used to monitor the activities of individual flies for 3 days on 12 hours light and 12 hours dark cycle. F1 male flies were collected immediately after eclosion, maintained in groups of 10 flies per vial and assayed at a desired age at 25°C. Flies were placed inside the activity monitor tubes for at least 24 hours before each experiment to acclimatize to the experimental set-up. 32 flies were placed in 5 mm diameter tubes, with one end containing 1 cm worth of food. *Drosophila* activity was recorded every 5 minutes. The raw data was processed using the actmon R package publicly available online at https://github.com/kazi11/actmon (kindly provided by Jeff Stafford). Data from flies that died during the experiment were discarded. The processed results were analysed using one-way ANOVA and Tukey’s multiple comparison tests using GraphPad Prism.

### Leg motoneuron analysis

To examine the impact of knockdown at a neuronal level, adult male *Drosophila* ventral nerve cords (VNCs) were dissected in ice cold 1x PBS, over a period of 30 minutes. The VNCs were fixed, on poly-Lysine coated slides, in 4% PFA at room temperature for 30 minutes followed by 3x 5 minutes washes with 0.1% PBT. They were mounted in Vectashield mounting media (Vector Laboratories Inc.), coverslipped and sealed with nail-polish. All images were acquired using the Fast Airyscan feature on a Zeiss LSM 880 inverted confocal microscope. Images were taken in a Z-stack and the leg motoneurons were quantified in a 3D-model (using Zen-blue image processing software) separately for T1, T2 and T3 segments. The results were analysed using one-way ANOVA and Tukey’s multiple comparison tests using GraphPad Prism.

---

**Supplementary Figure 1.**
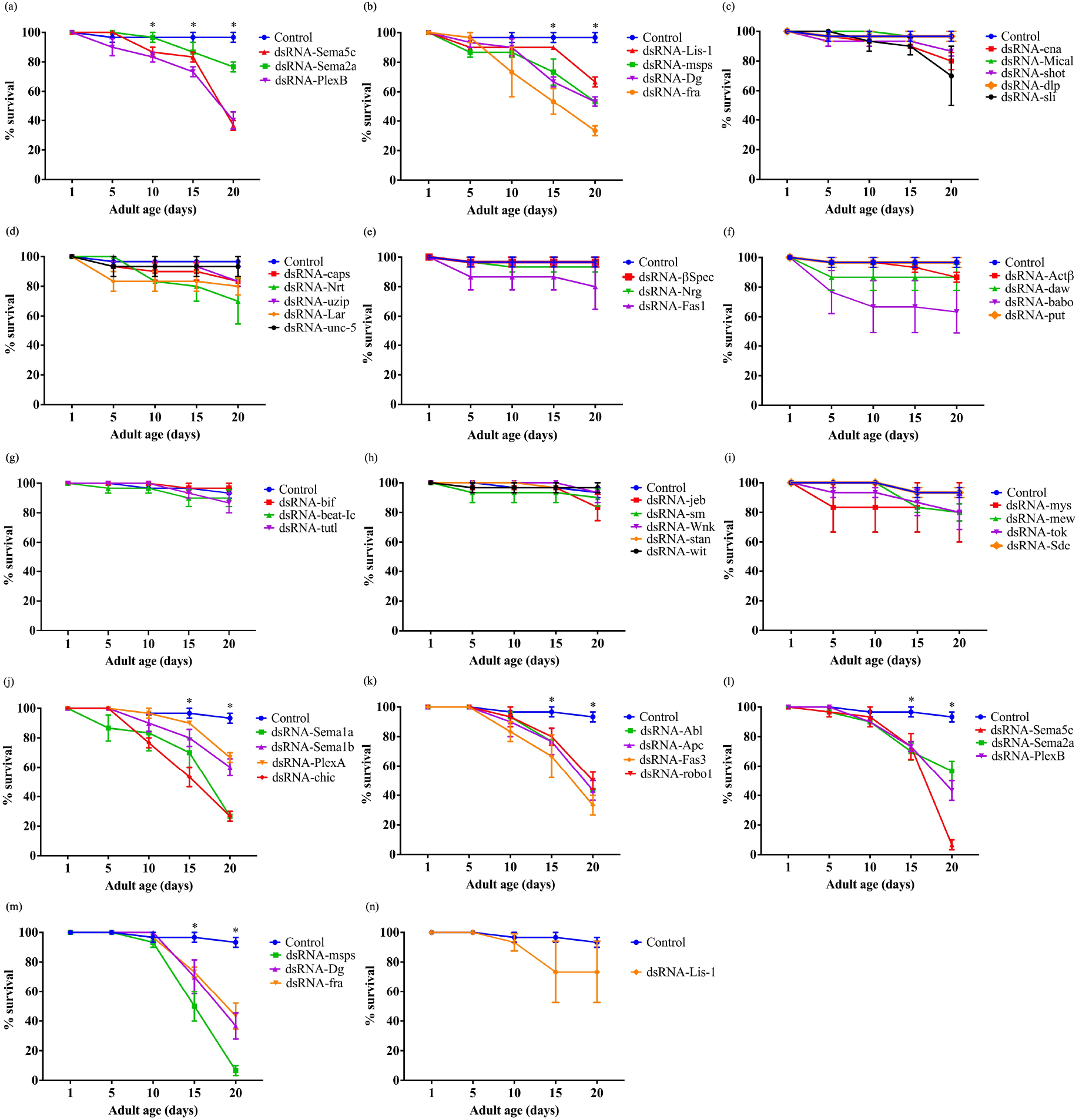
Knockdown of a subset of axon guidance genes reduces adult survival rate. (a)-(i) Survival analysis curves of additional genes from the GeneSwitch screen. (a) Genes that exhibited a significant effect on survival from day 9 post eclosion. (b) Genes that exhibited a significant effect from day 14. (c)- (i) Genes that exhibited no significant effect on adult survival rate until day 20. (j)-(n) 14 of the 15 axon guidance genes that resulted in a significant reduction in the adult survival rate in the GeneSwitch screen also reduced survival rate in the TARGET screen. Each data point is the average of three replicates. Data shown are mean ± SEM (p < 0.05, two-way ANOVA and Tukey’s multiple comparison tests). ‘*’ indicates the timepoints when the survival of knockdowns was significantly different from that of the controls.

**Supplementary Figure 2.**
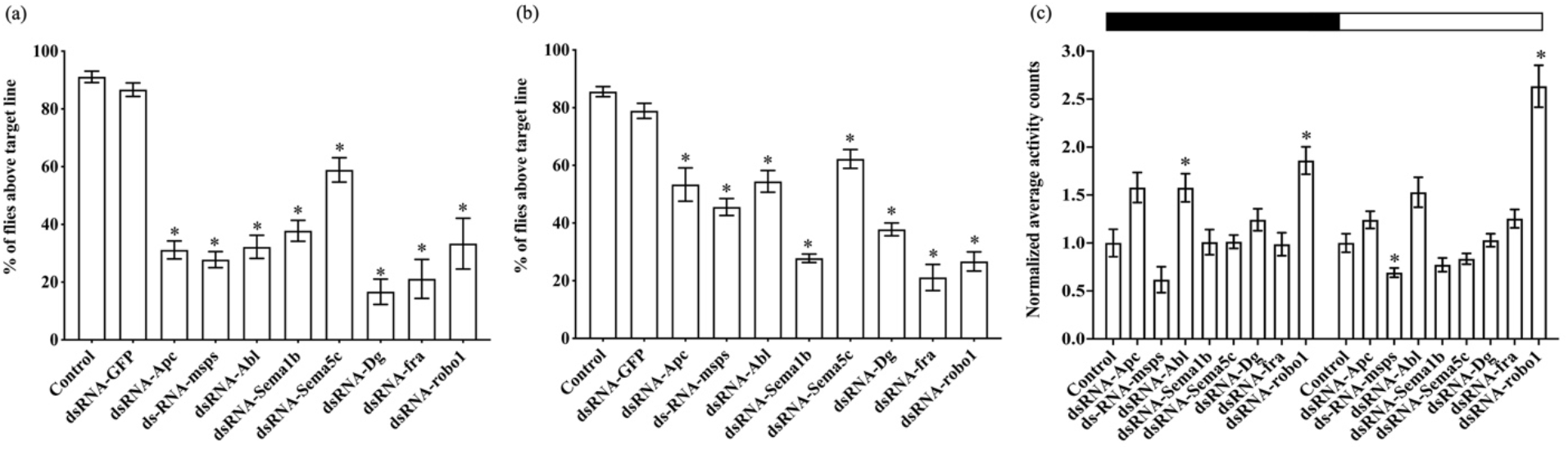
Motility is impacted by the knockdown of specific axon guidance genes. (a) Effect of pan-neuronal knockdown of *Apc, msps, Abl, Sema1b, Sema5c, Dg, fra* and *robo1* (using *elav-GeneSwitch-Gal4*) on adult climbing behavior. A significant decrease in the adult climbing ability was observed after knockdown. (b) Similar observations were made when each gene was knocked down specifically in glutamatergic motoneurons using the *OK371-Gal4*. Data shown are mean ± SEM. Each data point is the average of three replicates. (p < 0.05, oneway ANOVA and Tukey’s multiple comparison tests) (c) Effect of pan-neuronal knockdown of axon guidance genes (using *elav-GeneSwitch-Gal4*) on adult activity (measured using DAM). The bar graph compares the normalized average activity counts per 12 hours for each gene from multiple flies over a period of 3 days in 12 hours light and 12 hours dark cycle. The black portion (0-12 hours) represents the dark cycle and the white portion (12-24 hours) represents the light cycle. Data shown are mean ± SEM (‘*’ denotes p < 0.05, one-way ANOVA and Tukey’s multiple comparison tests).

**Supplementary Figure 3.**
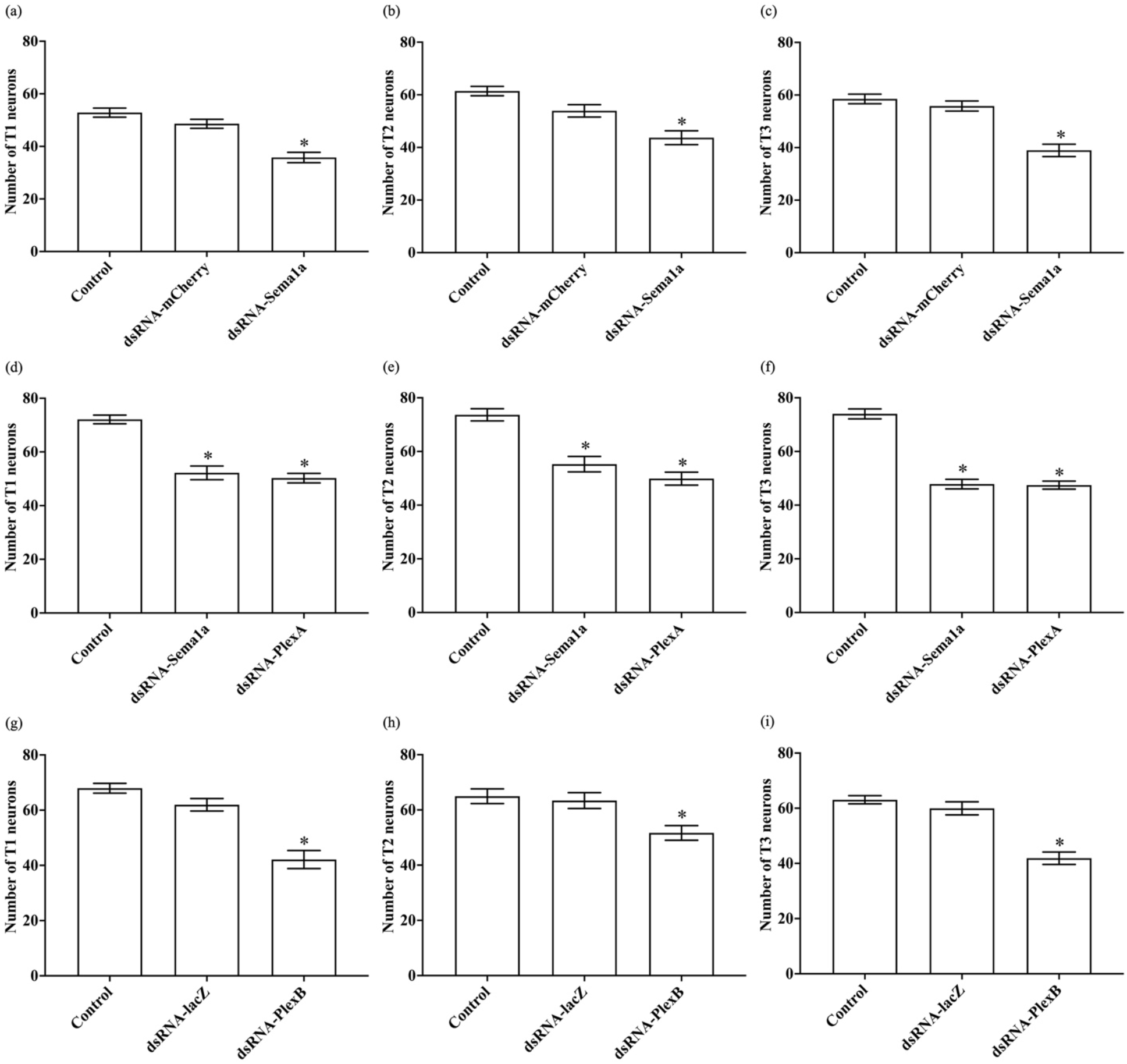
Knockdown of Semaphorins and Plexins results in motoneuron death. Quantification of adult leg motoneurons in T1 (a, d, g), T2 (b, e, h) and T3 (c, f, i) thoracic segments, following 14 days of axon guidance gene knockdown using additional *UAS-dsRNA* lines against Semaphorins and Plexins. Data shown are mean ± SEM (‘*’ denotes p < 0.05, one-way ANOVA and Tukey’s multiple comparison tests).

## References

1) Tessier-Lavigne, M., & Goodman, C. S. (1996). The molecular biology of axon guidance. Science, 274(5290), 1123–1133.

2) French, L., & Pavlidis, P. (2011). Relationships between gene expression and brain wiring in the adult rodent brain. PLoS Comput Biol, 7(1), e1001049.

3) Giger, R. J., Hollis, E. R., & Tuszynski, M. H. (2010). Guidance molecules in axon regeneration. Cold Spring Harbor perspectives in biology, 2(7), a001867.

4) Tanelian, D. L., Barry, M. A., Johnston, S. A., & Smith, G. M. (1997). Semaphorin III can repulse and inhibit adult sensory afferents in vivo. Nature medicine, 3(12), 1398–1401.

5) Kikuchi, K., Kishino, A., Konishi, O., Kumagai, K., Hosotani, N., Saji, I.,… & Kimura, T. (2003). In vitro and in vivo characterization of a novel semaphorin 3A inhibitor, SM-216289 or xanthofulvin. Journal of Biological Chemistry, 278(44), 42985–42991.

6) Pasterkamp, R. J., & Verhaagen, J. (2006). Semaphorins in axon regeneration: developmental guidance molecules gone wrong?. Philosophical Transactions of the Royal Society of London B:Biological Sciences, 361(1473), 1499–1511.

7) Fabes, J., Anderson, P., Brennan, C., & Bolsover, S. (2007). Regeneration-enhancing effects of EphA4 blocking peptide following corticospinal tract injury in adult rat spinal cord. European Journal of Neuroscience, 26(9), 2496–2505.

8) Horn, K. E., Glasgow, S. D., Gobert, D., Bull, S. J., Luk, T., Girgis, J.,… & Hamel, E. (2013). DCC expression by neurons regulates synaptic plasticity in the adult brain. Cell reports, 3(1), 173–185.

9) O’Connor, T. P., Cockburn, K., Wang, W., Tapia, L., Currie, E., & Bamji, S. X. (2009). Semaphorin 5B mediates synapse elimination in hippocampal neurons. Neural development, 4(1), 18.

10) Van Battum, E. Y., Brignani, S., & Pasterkamp, R. J. (2015). Axon guidance proteins in neurological disorders. The Lancet Neurology, 14(5), 532–546.

11) Graveley, B. R., Brooks, A. N., Carlson, J. W., Duff, M. O., Landolin, J. M., Yang, L.,… & Celniker, S. E. (2011). The developmental transcriptome of Drosophila melanogaster. Nature, 471(7339), 473–479.

12) Gramates, L. S., Agapite, J., Attrill, H., Calvi, B. R., Crosby, M. A., Dos Santos, G.,… & Strelets, V. B. (2022). FlyBase: A guided tour of highlighted features. Genetics, 220(4), iyac035.

13) Gelbart, W. M., & Emmert, D. B. (2013). Flybase high throughput expression pattern data. FlyBase Analysis (flybaseorg/reports/FBrf0221009html 29 October 20l3, date last accessed).

14) McGuire, S. E., Mao, Z., & Davis, R. L. (2004). Spatiotemporal gene expression targeting with the TARGET and gene-switch systems in Drosophila. Science’s STKE, 2004(220), p16–p16.

15) Zapala, M. A., Hovatta, I., Ellison, J. A., Wodicka, L., Del Rio, J. A., Tennant, R.,… & Winrow, C. (2005). Adult mouse brain gene expression patterns bear an embryologic imprint. Proceedings of the National Academy of Sciences of the United States of America, 102(29), 10357–10362.

16) Murray, K. D., Choudary, P. V., & Jones, E. G. (2007). Nucleus-and cell-specific gene expression in monkey thalamus. Proceedings of the National Academy of Sciences, 104(6), 1989–1994.

17) Hu, Y., Roesel, C., Flockhart, I., Perkins, L., Perrimon, N., & Mohr, S. E. (2013). UP-TORR: online tool for accurate and Up-to-Date annotation of RNAi Reagents. Genetics, 195(1), 37–45.

