## Supplementary Table 1 for "ADULT EXPRESSION OF SEMAPHORINS AND PLEXINS IS ESSENTIAL FOR MOTONEURON SURVIVAL"

**Supplementary Table 1. List of 162 genes categorized under the GO term ‘axon guidance’ that are expressed in the adult *Drosophila* nervous system*.***

| **#** | **Symbol** | **Name** | **Annotation ID** |
| --- | --- | --- | --- |
| 1 | 14-3-3ε | 14-3-3ε | CG31196 |
| 2 | aay | astray | CG3705 |
| 3 | Abl | Abl tyrosine kinase | CG4032 |
| 4 | acj6 | abnormal chemosensory jump 6 | CG9151 |
| 5 | Acsl | Acyl-CoA synthetase long-chain | CG8732 |
| 6 | ago | archipelago | CG15010 |
| 7 | Alk | Alk | CG8250 |
| 8 | aos | argos | CG4531 |
| 9 | AP-1σ | Adaptor Protein complex 1, σ subunit | CG5864 |
| 10 | ap | apterous | CG8376 |
| 11 | Apc | APC-like | CG1451 |
| 12 | Apc2 | Adenomatous polyposis coli tumor suppressor homolog 2 | CG6193 |
| 13 | babo | baboon | CG8224 |
| 14 | beat-Ia | beaten path Ia | CG4846 |
| 15 | beat-Ic | beaten path Ic | CG4838 |
| 16 | bif | bifocal | CG1822 |
| 17 | bon | bonus | CG5206 |
| 18 | brat | brain tumor | CG10719 |
| 19 | bsk | basket | CG5680 |
| 20 | bur | burgundy | CG9242 |
| 21 | CadN | Cadherin-N | CG7100 |
| 22 | caps | capricious | CG11282 |
| 23 | Cdc42 | Cdc42 | CG12530 |
| 24 | Cdk8 | Cyclin-dependent kinase 8 | CG10572 |
| 25 | CG4203 | - | CG4203 |
| 26 | chb | chromosome bows | CG32435 |
| 27 | Chi | Chip | CG3924 |
| 28 | chic | chickadee | CG9553 |
| 29 | CkIIα | casein kinase IIα | CG17520 |
| 30 | ckn | caskin | CG12424 |
| 31 | comm | commissureless | CG17943 |
| 32 | dac | dachshund | CG4952 |
| 33 | daw | dawdle | CG16987 |
| 34 | Dbx | Dbx | CG42234 |
| 35 | Dg | Dystroglycan | CG18250 |
| 36 | dlp | dally-like | CG32146 |
| 37 | dnt | doughnut on 2 | CG17559 |
| 38 | dock | dreadlocks | CG3727 |
| 39 | drl | derailed | CG17348 |
| 40 | Dscam1 | Down syndrome cell adhesion molecule 1 | CG17800 |
| 41 | E(z) | Enhancer of zeste | CG6502 |
| 42 | egh | egghead | CG9659 |
| 43 | en | engrailed | CG9015 |
| 44 | ena | enabled | CG15112 |
| 45 | Ephrin | Ephrin | CG1862 |
| 46 | Fas1 | Fasciclin 1 | CG6588 |
| 47 | Fas3 | Fasciclin 3 | CG5803 |
| 48 | Fmr1 | Fmr1 | CG6203 |
| 49 | Fps85D | Fps oncogene analog | CG8874 |
| 50 | fra | frazzled | CG8581 |
| 51 | gogo | golden goal | CG32227 |
| 52 | gt | giant | CG7952 |
| 53 | gukh | GUK-holder | CG31043 |
| 54 | Hr51 | Hormone receptor 51 | CG16801 |
| 55 | Hrb27C | Heterogeneous nuclear ribonucleoprotein at 27C | CG10377 |
| 56 | Hsc70-4 | Heat shock protein cognate 4 | CG4264 |
| 57 | hts | hu li tai shao | CG43443 |
| 58 | if | inflated | CG9623 |
| 59 | InR | Insulin-like receptor | CG18402 |
| 60 | jeb | jelly belly | CG30040 |
| 61 | jing | jing | CG9397 |
| 62 | Khc | Kinesin heavy chain | CG7765 |
| 63 | kis | kismet | CG3696 |
| 64 | Klp64D | Kinesin-like protein at 64D | CG10642 |
| 65 | Kr | Kruppel | CG3340 |
| 66 | kuz | kuzbanian | CG7147 |
| 67 | LanA | Laminin A | CG10236 |
| 68 | Lar | Leukocyte-antigen-related-like | CG10443 |
| 69 | lea | leak | CG5481 |
| 70 | Liprin-α | Liprin-α | CG11199 |
| 71 | lola | longitudinals lacking | CG12052 |
| 72 | meigo | medial glomeruli | CG5802 |
| 73 | mew | multiple edematous wings | CG1771 |
| 74 | Mical | Molecule interacting with CasL | CG33208 |
| 75 | mid | midline | CG6634 |
| 76 | msn | misshapen | CG16973 |
| 77 | msps | mini spindles | CG5000 |
| 78 | mys | myospheroid | CG1560 |
| 79 | N | Notch | CG3936 |
| 80 | Nedd4 | Nedd4 | CG42279 |
| 81 | nerfin-1 | nervous fingers 1 | CG13906 |
| 82 | NetA | Netrin-A | CG18657 |
| 83 | NetB | Netrin-B | CG10521 |
| 84 | Nf-YC | Nuclear factor Y-box C | CG3075 |
| 85 | NijA | Ninjurin A | CG6449 |
| 86 | NiPp1 | Nuclear inhibitor of Protein phosphatase 1 | CG8980 |
| 87 | not | non-stop | CG4166 |
| 88 | Nrt | Neurotactin | CG9704 |
| 89 | NT1 | Neurotrophin 1 | CG42576 |
| 90 | nvy | nervy | CG3385 |
| 91 | otk | off-track | CG8967 |
| 92 | Pak | PAK-kinase | CG10295 |
| 93 | pdm3 | pou domain motif 3 | CG42698 |
| 94 | Pka-R2 | cAMP-dependent protein kinase R2 | CG15862 |
| 95 | plexA | plexin A | CG11081 |
| 96 | plexB | plexin B | CG17245 |
| 97 | pod1 | pod1 | CG4532 |
| 98 | Pp1-87B | Protein phosphatase 1 at 87B | CG5650 |
| 99 | pros | prospero | CG17228 |
| 100 | Psc | Posterior sex combs | CG3886 |
| 101 | ptc | patched | CG2411 |
| 102 | Ptp52F | Ptp52F | CG18243 |
| 103 | Ptp61F | Protein tyrosine phosphatase 61F | CG9181 |
| 104 | Ptp69D | Protein tyrosine phosphatase 69D | CG10975 |
| 105 | put | punt | CG7904 |
| 106 | Rab6 | Rab6 | CG6601 |
| 107 | Rac1 | Rac1 | CG2248 |
| 108 | raptor | raptor | CG4320 |
| 109 | ras | raspberry | CG1799 |
| 110 | retn | retained | CG5403 |
| 111 | Rheb | Ras homolog enriched in brain ortholog (H. sapiens) | CG1081 |
| 112 | Rho1 | Rho1 | CG8416 |
| 113 | RhoGAP93B | Rho GTPase activating protein at 93B | CG3421 |
| 114 | RhoGEF64C | Rho guanine nucleotide exchange factor at 64C | CG32239 |
| 115 | Rich | RIC1 homolog | CG9063 |
| 116 | robo | roundabout | CG13521 |
| 117 | robo3 | robo3 | CG5423 |
| 118 | run | runt | CG1849 |
| 119 | S6k | RPS6-p70-protein kinase | CG10539 |
| 120 | sbb | scribbler | CG5580 |
| 121 | scb | scab | CG8095 |
| 122 | Sdc | Syndecan | CG10497 |
| 123 | sec15 | sec15 | CG7034 |
| 124 | Sema-1a | Sema-1a | CG18405 |
| 125 | Sema-1b | Sema-1b | CG6446 |
| 126 | Sema-2a | Sema-2a | CG4700 |
| 127 | Sema-2b | Semaphorin-2b | CG33960 |
| 128 | Sema-5c | Semaphorin-5c | CG5661 |
| 129 | sens | senseless | CG32120 |
| 130 | seq | sequoia | CG32904 |
| 131 | shg | shotgun | CG3722 |
| 132 | sim | single-minded | CG7771 |
| 133 | sli | slit | CG43758 |
| 134 | sm | smooth | CG9218 |
| 135 | Smox | Smad on X | CG2262 |
| 136 | SNF1A | SNF1A/AMP-activated protein kinase | CG3051 |
| 137 | SoxN | SoxNeuro | CG18024 |
| 138 | spen | split ends | CG18497 |
| 139 | spz5 | spatzle 5 | CG9972 |
| 140 | Sra-1 | specifically Rac1-associated protein 1 | CG4931 |
| 141 | stan | starry night | CG11895 |
| 142 | Tig | Tiggrin | CG11527 |
| 143 | tok | tolkin | CG6863 |
| 144 | Tor | Target of rapamycin | CG5092 |
| 145 | Trim9 | Trim9 | CG31721 |
| 146 | trio | trio | CG18214 |
| 147 | trx | trithorax | CG8651 |
| 148 | ttv | tout-velu | CG10117 |
| 149 | tup | tailup | CG10619 |
| 150 | tutl | turtle | CG15427 |
| 151 | unc-5 | unc-5 | CG8166 |
| 152 | unc-104 | unc-104 ortholog | CG8566 |
| 153 | Unc-115a | - | CG31352 |
| 154 | uzip | unzipped | CG3533 |
| 155 | Vav | Vav ortholog (H. sapiens) | CG7893 |
| 156 | velo | veloren | CG10107 |
| 157 | Wnk | WNK homolog | CG7177 |
| 158 | Wnt5 | Wnt oncogene analog 5 | CG6407 |
| 159 | Xe7 | Xe7 | CG2179 |
| 160 | XNP | XNP | CG4548 |
| 161 | β-Spec | β Spectrin | CG5870 |
| 162 | βTub60D | β-Tubulin at 60D | CG3401 |
