## Supplementary Table 2 for "ADULT EXPRESSION OF SEMAPHORINS AND PLEXINS IS ESSENTIAL FOR MOTONEURON SURVIVAL"

**Supplementary Table 2. List of 122 genes categorized under the GO term ‘dendrite morphogenesis’ that are expressed in the adult *Drosophila* nervous system*.***

| **#** | **Symbol** | **Name** | **Annotation ID** |
| --- | --- | --- | --- |
| 1 | Aats-gln | Glutaminyl-tRNA synthetase | CG10506 |
| 2 | Aats-gly | Glycyl-tRNA synthetase | CG6778 |
| 3 | Aats-trp | Tryptophanyl-tRNA synthetase | CG9735 |
| 4 | ab | abrupt | CG43860 |
| 5 | Acf1 | ATP-dependent chromatin assembly factor large subunit | CG1966 |
| 6 | acj6 | abnormal chemosensory jump 6 | CG9151 |
| 7 | Actβ | Activin-β | CG11062 |
| 8 | Adf1 | Adh transcription factor 1 | CG15845 |
| 9 | aop | anterior open | CG3166 |
| 10 | Arc42 | Arc42 | CG4703 |
| 11 | Ark | Apaf-1-related-killer | CG6829 |
| 12 | asf1 | anti-silencing factor 1 | CG9383 |
| 13 | babo | baboon | CG8224 |
| 14 | Bap55 | Brahma associated protein 55kD | CG6546 |
| 15 | Bap60 | Brahma associated protein 60kD | CG4303 |
| 16 | bigmax | bigmax | CG3350 |
| 17 | bon | bonus | CG5206 |
| 18 | brm | brahma | CG5942 |
| 19 | Caf1 | Chromatin assembly factor 1 subunit | CG4236 |
| 20 | Cdc42 | Cdc42 | CG12530 |
| 21 | CG2678 | - | CG2678 |
| 22 | CG4328 | - | CG4328 |
| 23 | CG5343 | - | CG5343 |
| 24 | CG9104 | - | CG9104 |
| 25 | CG34340 | - | CG34340 |
| 26 | chinmo | Chronologically inappropriate morphogenesis | CG31666 |
| 27 | chm | chameau | CG5229 |
| 28 | ci | cubitus interruptus | CG2125 |
| 29 | comm | commissureless | CG17943 |
| 30 | Crg-1 | Circadianly Regulated Gene | CG32788 |
| 31 | ct | cut | CG11387 |
| 32 | Cul-3 | Cullin-3 | CG42616 |
| 33 | d4 | d4 | CG2682 |
| 34 | dally | division abnormally delayed | CG4974 |
| 35 | Dcr-1 | Dicer-1 | CG4792 |
| 36 | Dhc64C | Dynein heavy chain 64C | CG7507 |
| 37 | Dlic | Dynein light intermediate chain | CG1938 |
| 38 | E(bx) | Enhancer of bithorax | CG32346 |
| 39 | E(z) | Enhancer of zeste | CG6502 |
| 40 | E2f | E2F transcription factor | CG6376 |
| 41 | EcR | Ecdysone receptor | CG1765 |
| 42 | Elongin-C | Elongin C | CG9291 |
| 43 | ems | empty spiracles | CG2988 |
| 44 | ena | enabled | CG15112 |
| 45 | esc | extra sexcombs | CG14941 |
| 46 | Fmr1 | Fmr1 | CG6203 |
| 47 | fra | frazzled | CG8581 |
| 48 | fru | fruitless | CG14307 |
| 49 | futsch | futsch | CG34387 |
| 50 | Gcn5 | Gcn5 ortholog | CG4107 |
| 51 | gro | groucho | CG8384 |
| 52 | ham | hamlet | CG31753 |
| 53 | HmgD | High mobility group protein D | CG17950 |
| 54 | Iswi | Imitation SWI | CG8625 |
| 55 | jumu | jumeau | CG4029 |
| 56 | Khc | Kinesin heavy chain | CG7765 |
| 57 | kn | knot | CG10197 |
| 58 | kni | knirps | CG4717 |
| 59 | l(3)mbt | lethal (3) malignant brain tumor | CG5954 |
| 60 | Lis-1 | Lissencephaly-1 | CG8440 |
| 61 | mbf1 | multiprotein bridging factor 1 | CG4143 |
| 62 | MED4 | Mediator complex subunit 4 | CG8609 |
| 63 | MED11 | Mediator complex subunit 11 | CG6884 |
| 64 | MED24 | Mediator complex subunit 24 | CG7999 |
| 65 | MEP-1 | - | CG1244 |
| 66 | Mi-2 | - | CG8103 |
| 67 | Mical | Molecule interacting with CasL | CG33208 |
| 68 | Nak | Numb-associated kinase | CG10637 |
| 69 | Nc | Nedd2-like caspase | CG8091 |
| 70 | Not1 | Not1 | CG34407 |
| 71 | Nrg | Neuroglian | CG1634 |
| 72 | nvy | nervy | CG3385 |
| 73 | Pi3K92E | Pi3K92E | CG4141 |
| 74 | prel | preli-like | CG8806 |
| 75 | pros | prospero | CG17228 |
| 76 | Ptp69D | Protein tyrosine phosphatase 69D | CG10975 |
| 77 | Ptx1 | Ptx1 | CG1447 |
| 78 | pum | pumilio | CG9755 |
| 79 | put | punt | CG7904 |
| 80 | pygo | pygopus | CG11518 |
| 81 | Rab5 | Rab5 | CG3664 |
| 82 | Rac1 | Rac1 | CG2248 |
| 83 | Rfx | Rfx | CG6312 |
| 84 | Rho1 | Rho1 | CG8416 |
| 85 | rictor | rapamycin-insensitive companion of Tor | CG8002 |
| 86 | robl | roadblock | CG10751 |
| 87 | robo | roundabout | CG13521 |
| 88 | Rpd3 | Rpd3 | CG7471 |
| 89 | run | runt | CG1849 |
| 90 | S6k | RPS6-p70-protein kinase | CG10539 |
| 91 | scrt | scratch | CG1130 |
| 92 | Sema-1a | Sema-1a | CG18405 |
| 93 | seq | sequoia | CG32904 |
| 94 | shot | short stop | CG18076 |
| 95 | shrb | shrub | CG8055 |
| 96 | Sin1 | SAPK-interacting protein 1 | CG10105 |
| 97 | Sin3A | Sin3A | CG8815 |
| 98 | Sirt2 | Sirt2 | CG5085 |
| 99 | sli | slit | CG43758 |
| 100 | SMC1 | SMC1 | CG6057 |
| 101 | Smox | Smad on X | CG2262 |
| 102 | Snr1 | Snf5-related 1 | CG1064 |
| 103 | Sox14 | Sox box protein 14 | CG3090 |
| 104 | sqz | squeeze | CG5557 |
| 105 | stan | starry night | CG11895 |
| 106 | Su(z)12 | Su(z)12 | CG8013 |
| 107 | sv | shaven | CG11049 |
| 108 | Tab2 | TAK1-associated binding protein 2 | CG7417 |
| 109 | Taf4 | TBP-associated factor 4 | CG5444 |
| 110 | Tango10 | Transport and Golgi organization 10 | CG1841 |
| 111 | TER94 | TER94 | CG2331 |
| 112 | tgo | tango | CG11987 |
| 113 | Tm1 | Tropomyosin 1 | CG4898 |
| 114 | Tor | Target of rapamycin | CG5092 |
| 115 | trc | tricornered | CG8637 |
| 116 | trh | trachealess | CG42865 |
| 117 | ttk | tramtrack | CG1856 |
| 118 | Ube3a | Ubiquitin protein ligase E3A | CG6190 |
| 119 | usp | ultraspiracle | CG4380 |
| 120 | vvl | ventral veins lacking | CG10037 |
| 121 | W | Wrinkled | CG5123 |
| 122 | wit | wishful thinking | CG10776 |
