## Supplementary Table 3 for "ADULT EXPRESSION OF SEMAPHORINS AND PLEXINS IS ESSENTIAL FOR MOTONEURON SURVIVAL"

**Supplementary Table 3. Expression levels of genes categorized under the GO term ‘axon guidance’ (excluding transcription factors) in the adult *Drosophila* nervous system*.* Yellow highlighted genes were prioritized for the screen. (RPKM- Reads Per Kilobase of transcript per Million reads mapped)**

| **Very low**  **(1- 3 RPKM)** | **Low**  **(4-10 RPKM)** | **Moderate**  **(11-25 RPKM)** | **Moderately high**  **(26-50 RPKM)** |
| --- | --- | --- | --- |
| plexin B | tout-velu | Leukocyte-antigen-related-like | astray |
| doughnut on 2 | Laminin A | Kinesin-like protein at 64D | Acyl-CoA synthetase long-chain |
| multiple edematous wings | Netrin-B | Ras homolog enriched in brain ortholog (H. sapiens) | Adaptor Protein complex 1, σ subunit |
| Ptp52F | brain tumor | plexin A | bifocal |
| Netrin-A | Protein tyrosine phosphatase 69D | capricious | burgundy |
| patched | Liprin-α | APC-like | chickadee |
| Trim9 | starry night | enabled | dawdle |
| golden goal | caskin | misshapen | egghead |
| beaten path Ic | roundabout | trio | Fasciclin 1 |
| beaten path Ia | archipelago | Dystroglycan | Fmr1 |
| leak | derailed | Sema-1a | Heterogeneous nuclear ribonucleoprotein at 27C |
| Semaphorin-5c | Down syndrome cell adhesion molecule 1 | jelly belly | Kinesin heavy chain |
| sec15 | commissureless | Unc-115a | myospheroid |
| scab | Insulin-like receptor | Molecule interacting with CasL | PAK-kinase |
| frazzled | Ephrin | β-Tubulin at 60D | cAMP-dependent protein kinase R2 |
| spatzle 5 | GUK-holder | Rho GTPase activating protein at 93B | Protein phosphatase 1 at 87B |
|  | dally-like | dreadlocks | Rac1 |
|  | Rho guanine nucleotide exchange factor at 64C | Abl tyrosine kinase | raspberry |
|  | Semaphorin-2b | hu li tai shao | Rho1 |
|  | shotgun | slit | RPS6-p70-protein kinase |
|  | CG4203 | pod1 | Syndecan |
|  | raptor | Sema-2a | Smad on X |
|  | argos | mini spindles | Tiggrin |
|  | specifically Rac1-associated protein 1 | Fasciclin 3 | tolkin |
|  | Target of rapamycin | Sema-1b | turtle |
|  | robo3 | Rab6 | unc-104 ortholog |
|  | medial glomeruli | WNK homolog | unzipped |
|  | Wnt oncogene analog 5 | baboon | Xe7 |
|  | Ninjurin A | Alk | β Spectrin |
|  | Cadherin-N | Fps oncogene analog |  |
|  | kuzbanian | smooth |  |
|  | Vav ortholog (H. sapiens) |  |  |
|  | punt |  |  |
|  | unc-5 |  |  |
|  | RIC1 homolog |  |  |
|  | inflated |  |  |
|  | Neurotactin |  |  |
