## Supplementary Table 4 for "ADULT EXPRESSION OF SEMAPHORINS AND PLEXINS IS ESSENTIAL FOR MOTONEURON SURVIVAL"

**Supplementary Table 4. Expression levels of genes categorized under the GO term ‘dendrite morphogenesis’ (excluding transcription factors) in the adult *Drosophila* nervous system*.* Yellow highlighted genes were prioritized for the screen. (RPKM- Reads Per Kilobase of transcript per Million reads mapped)**

| **Very low**  **(1- 3 RPKM)** | **Low**  **(4-10 RPKM)** |
| --- | --- |
| Wrinkled | Glutaminyl-tRNA synthetase |
| Nedd2-like caspase | Glycyl-tRNA synthetase |
| frazzled | Tryptophanyl-tRNA synthetase |
|  | Activin-β |
|  | Arc42 |
|  | Apaf-1-related-killer |
|  | anti-silencing factor 1 |
|  | baboon |
|  | Brahma associated protein 55kD |
|  | CG9104 |
|  | Chronologically inappropriate morphogenesis |
|  | commissureless |
|  | Cullin-3 |
|  | division abnormally delayed |
|  | Dynein heavy chain 64C |
|  | Dynein light intermediate chain |
|  | Enhancer of bithorax |
|  | Elongin C |
|  | enabled |
|  | Fmr1 |
|  | futsch |
|  | High mobility group protein D |
|  | Kinesin heavy chain |
|  | Lissencephaly-1 |
|  | Molecule interacting with CasL |
|  | Numb-associated kinase |
|  | Not1 |
|  | Neuroglian |
|  | Pi3K92E |
|  | preli-like |
|  | Protein tyrosine phosphatase 69D |
|  | pumilio |
|  | punt |
|  | Rab5 |
|  | Rac1 |
|  | Rho1 |
|  | rapamycin-insensitive companion of Tor |
|  | roadblock |
|  | roundabout |
|  | RPS6-p70-protein kinase |
|  | Sema-1a |
|  | short stop |
|  | shrub |
|  | SAPK-interacting protein 1 |
|  | Sirt2 |
|  | slit |
|  | Smad on X |
|  | starry night |
|  | Transport and Golgi organization 10 |
|  | TER94 |
|  | Tropomyosin 1 |
|  | Target of rapamycin |
|  | wishful thinking |
